# Neurometabolic timecourse of healthy aging

**DOI:** 10.1101/2022.06.08.495050

**Authors:** Tao Gong, Steve C.N. Hui, Helge J. Zöllner, Mark Britton, Yulu Song, Yufan Chen, Aaron T. Gudmundson, Kathleen E. Hupfeld, Saipavitra Murali-Manohar, Eric C. Porges, Georg Oeltzschner, Weibo Chen, Guangbin Wang, Richard A. E. Edden

**Affiliations:** Departments of Radiology, Shandong Provincial Hospital Affiliated to Shandong First Medical University, Jinan, Shandong, 250021, China; Departments of Radiology, Shandong Provincial Hospital, Shandong University, Jinan, Shandong, 250021, China; The Russell H. Morgan Department of Radiology and Radiological Science, Johns Hopkins University School of Medicine, Baltimore, MD, USA; F.M. Kirby Research Center for Functional Brain Imaging, Kennedy Krieger Institute, Baltimore, MD, USA; Center for Cognitive Aging and Memory, University of Florida, Gainesville, FL, USA; McKnight Brain Research Foundation, University of Florida, FL, USA; Department of Clinical and Health Psychology, University of Florida, Gainesville, FL, USA; Department of Neurobiology and Behavior, University of California, Irvine, CA, USA; Philips Healthcare, Shanghai, China

**Keywords:** magnetic resonance spectroscopy, neurometabolite, PRESS, healthy aging

## Abstract

**Purpose:** The neurometabolic timecourse of healthy aging is not well-established, in part due to diversity of quantification methodology. In this study, a large structured cross-sectional cohort of male and female subjects throughout adulthood was recruited to investigate neurometabolic changes as a function of age, using consensus-recommended magnetic resonance spectroscopy quantification methods.

**Methods:** 102 healthy volunteers, with approximately equal numbers of male and female participants in each decade of age from the 20s, 30s, 40s, 50s, and 60s, were recruited with IRB approval. MR spectroscopic data were acquired on a 3T MRI scanner. Metabolite spectra were acquired using PRESS localization (TE = 30 ms; 96 transients) in the centrum semiovale (CSO) and posterior cingulate cortex (PCC). Water-suppressed spectra were modeled using the Osprey algorithm, employing a basis set of 18 simulated metabolite basis functions and a cohort-mean measured macromolecular spectrum. Pearson correlations were conducted to assess relationships between metabolite concentrations and age for each voxel; paired t-tests were run to determine whether metabolite concentrations differed between the PCC and CSO.

**Results:** Two datasets were excluded (1 ethanol; 1 unacceptably large lipid signal). Statistically significant age-by-metabolite correlations were seen for tCr (R^2^=0.36; p<0.001), tCho (R^2^=0.11; p<0.001), sI (R^2^=0.11; p=0.004), and mI (R^2^=0.10; p<0.001) in the CSO, and tCr (R^2^=0.15; p<0.001), tCho (R^2^=0.11; p<0.001), and GABA (R^2^=0.11; p=0.003) in the PCC. No significant correlations were seen between tNAA, NAA, GSH, Glx or Glu and age in either region (all p>0.25). Levels of sI were significantly higher in the PCC in female subjects (p<0.001) than in male subjects. There was a significant positive correlation between linewidth and age.

**Conclusion:** The results indicated age correlations for tCho, tCr, sI, and mI in CSO and for tCr, tCho and GABA in PCC, while no age-related changes were found for NAA, tNAA, GSH, Glu or Glx. Our results provide a normative foundation for future work investigating the neurometabolic time course of healthy aging using MRS.

**Highlights:**

1. A large structured cross-sectional cohort of neurometabolic aging dataset is presented;
2. Age correlations were observed for tCho, tCr, sI, and mI in CSO and for tCr, tCho and GABA in PCC;
3. No age correlations were found for NAA, tNAA, GSH, Glu or Glx in either region.

## Introduction

With the global population aging and the prevalence of Alzheimer’s disease increasing (Langa et al., 2004), the study of the biochemical mechanisms of healthy and pathological aging is a major research priority. While cell-level neuroscience offers maximum scientific control and analytic precision, the need to link neurometabolic changes in the brain to changes in cognition, especially with a view to developing neuroprotective interventions, demands *in vivo* imaging methods. In-vivo magnetic resonance spectroscopy (MRS) of the brain can potentially bridge between cellular neuroscience and in vivo imaging of physical properties of tissue water by measuring the concentration of endogenous metabolites, particularly those associated with neurotransmission, energy metabolism and oxidative stress defense.

Major neurometabolites quantifiable by MRS include the neuronal marker N-acetyl aspartate (NAA) (Landim et al., 2016), as well as γ-aminobutyric acid (GABA) (Mullins et al., 2014), the major inhibitory neurotransmitter within the brain, and glutamate (Glu) (Cheng et al., 2021), the principal excitatory neurotransmitter. Glutamine (Gln) is an MRS-detectable precursor for Glu, though often MRS studies performed at 3T report Glx, the combination of Glu + Gln signals. N-acetyl aspartyl glutamate (NAAG) functions as a neuromodulator, inhibiting synaptic release of GABA, Glu, and dopamine (Harris et al., 2017). Aspartate (Asp) is an excitatory neuromodulator (Menshchikov et al., 2017) and precursor of NAA. Myo-Inositol (mI) acts as an osmolyte, with involvement in maintaining cell volume and fluid balance (Dai et al., 2016) as well as brain cell signaling (Hoyer et al., 2014) and glial cell proliferation (Brand et al., 1993). Scyllo-Inositol (sI) is formed from mI; the functional role of sI in the brain is less clear, though it may decrease accumulation of amyloid-beta protein (McLaurin et al., 2000). The overlapping choline signals (tCho) from free choline, glycerophosphocholine (GPC) and phosphocholine (PCh) are a cell membrane marker which reflects changes in membrane turnover or cell density (Cleeland et al., 2019). Creatine and phosphocreatine (reported in combination as tCr) and lactate (Lac) are all involved in energy metabolism. Creatine is a brain osmolyte and involved in maintenance of brain energy homeostasis (Ross and Sachdev, 2004). Lac is the end product of anaerobic glycolysis, and is found in very low concentrations in the brain under normal physiologic conditions (Harris et al., 2017), but elevated in conditions of altered energy metabolism such as tumor or stroke (Howe et al., 2003; Morana et al., 2015). Glutathione (GSH) is one of the most abundant antioxidant sources in the central nervous system and plays a key role in the maintenance of redox homeostasis (Dwivedi et al., 2020).

A number of cross-sectional studies have characterized the neurometabolic trajectory of aging. Although the results were varied, the most consistent findings demonstrated that NAA and Glu concentration decrease with age, while Cho, Cr and mI concentration increase with age (Cleeland et al., 2019; Haga et al., 2009). Studies using edited MRS methods targeting specific metabolites often show age-related decrease in GABA levels (Gao et al., 2013; Porges et al., 2021) which may be driven by tissue changes (Maes et al., 2018; Porges et al., 2017), and have demonstrated an age-related increase in GSH (Hupfeld et al., 2021).

This substantial body of MRS-aging literature has employed diverse methodological approaches, in terms of study design, acquisition, and quantification (Cleeland et al., 2019; Haga et al., 2009). A majority of studies (∼65%) has used a dichotomized young-old between-groups design, and of those studies that do consider age as a continuous variable, several have bimodal age distributions. The median total cohort size is 62 subjects. In terms of acquisition, a majority of studies (∼45%) has used PRESS localization for single-voxel acquisitions, whereas some (∼30%) studies employed multi-voxel MRSI methods. The quantification approaches range from metabolite ratios (∼30%) through water-referenced concentrations with CSF correction (∼30%) to full tissue-corrected water referencing (∼10%). Given the relatively diverse findings of this literature, which derive in part from limited statistical power, low SNR and methodological diversity, we designed a large structured cross-sectional cohort of male and female subjects throughout adulthood to investigate neurometabolic changes as a function of age, and used data processing, modeling, and quantification practices recommended by recent MRS expert community consensus (Near et al., 2021).

## Methods

### Participants

One hundred and two healthy volunteers were recruited with local IRB approval (Shandong Provincial Hospital). The cohort was structured to include approximately equal numbers of male and female participants in each decade of age from the 20s, 30s, 40s, 50s, to the 60s. Exclusion criteria included contraindications for MRI and a history of neurological and psychiatric illness. Metabolite-nulled data from the same cohort of subjects was recently published (Hui et al., 2022a) to investigate the age trajectory of macromolecular signals in the spectrum. Since this analysis revealed no significant age- or sex-related changes to the macromolecular spectrum, a cohort-mean macromolecule spectrum was incorporated into the modeling (see *Analysis*).

### MR protocol

Data were acquired on a 3T MRI scanner (Ingenia CX, Philips Healthcare, The Netherlands). Acquisition of MRS data was preceded by a T_1_-weighted MPRAGE scan (TR/TE/ 6.9/3.2 ms; FA 8°) with 1 mm^3^ isotropic resolution for voxel positioning and tissue segmentation. Metabolite spectra were acquired using PRESS localization (1.3 kHz refocusing bandwidth) with the following parameters: TR/TE: 2000/30 ms; 30 × 26 × 26 mm^3^ voxels localized in the CSO (predominantly white matter) and PCC (predominantly gray matter), as shown in Figure 1; 96 transients sampled at 2 kHz; water suppression was performed using the VAPOR method (Tkác et al., 1999). A slice-selective saturation pulse (20 mm thickness) was applied to suppress subcutaneous lipid adjacent to the voxel in CSO and PCC acquisitions. Water reference spectra were acquired without water suppression or pre-inversion.

**Figure 1.**
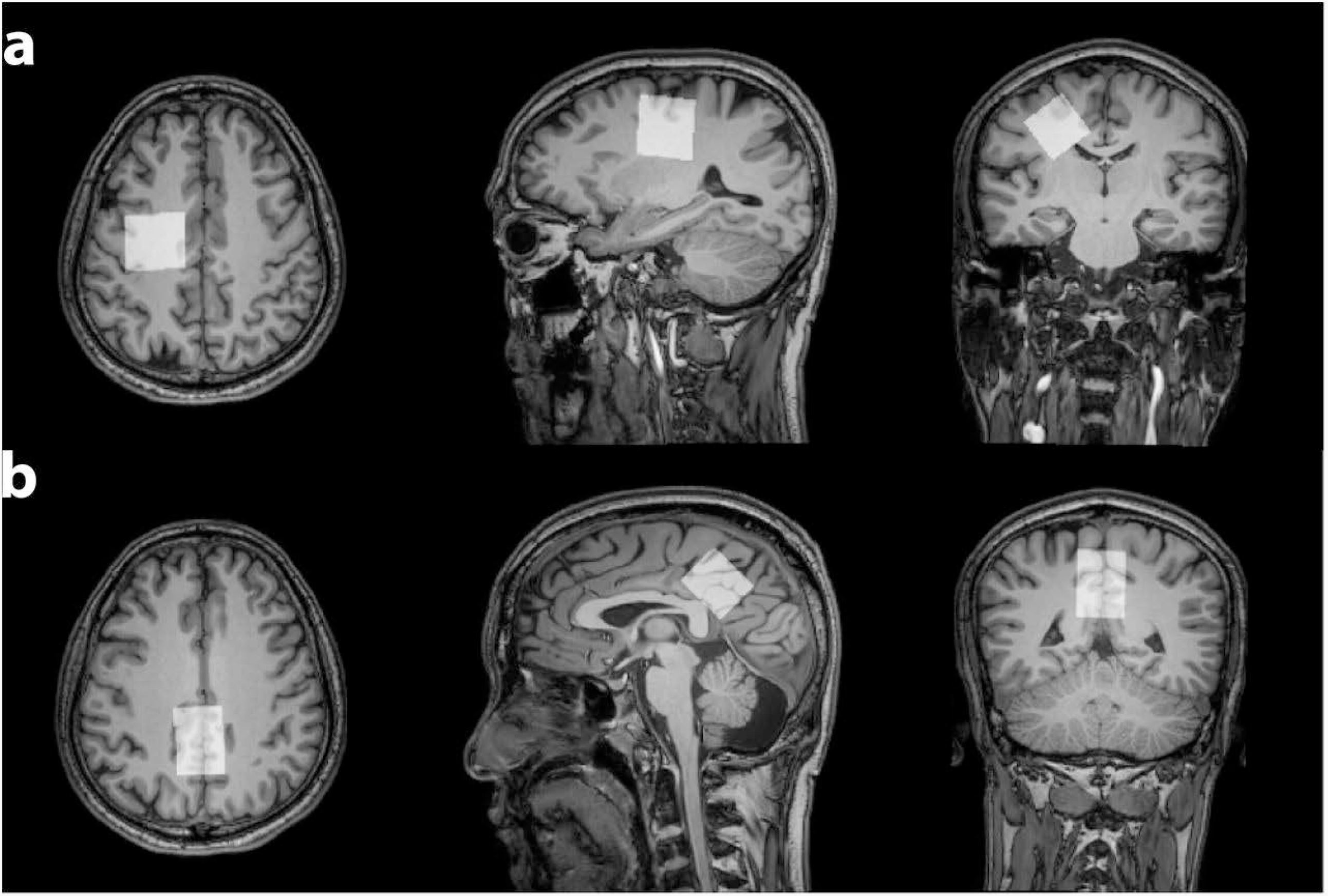
Voxels of interest a) left CSO and b) midline PCC in which spectral data were acquired. Acquisition parameters: 96 transients; TR = 2s; TE = 30 ms; 20 cm^3^ voxels.

### Analysis

T_1_-weighted images were segmented using SPM12 (Friston et al., 1994) algorithms called within Osprey (Oeltzschner et al., 2020) after voxel co-registration. Water-suppressed spectra were modeled using the Osprey algorithm, employing a basis set consisting of 18 simulated metabolite basis functions which were generated from a fully localized 2D density-matrix simulation of a 101 × 101 spatial grid (field of view 50% larger than voxel) using real pulse waveforms and sequence timings, as implemented in a MATLAB-based simulation toolbox FID-A (Simpson et al., 2017). Metabolites included in the model are as follows: ascorbate, Asp; creatine, Cr; negative creatine methylene, CrCH2; gamma-aminobutyric acid, GABA; glycerophosphocholine, GPC; glutathione, GSH; glutamine, Gln; glutamate, Glu; myo-inositol, mI; lactate, Lac; N-acetylaspartate, NAA; N-acetylaspartylglutamate, NAAG; phosphocholine, PCh; phosphocreatine, PCr; phosphoethanolamine, PE; scyllo-inositol, sI; and taurine, Tau. Osprey analysis procedures match those previously described in, with the exception that experimentally derived in vivo macromolecular (MM) basis spectra derived from a previous study (Hui et al., 2022b) are incorporated into the basis set instead of 8 parameterized Gaussian basis functions. To create the MM basis function, individual-subject ‘clean’ MM spectra (separate for PCC and CSO) were modeled with a flexible spline (0.1 ppm knot spacing) across the full spectral range. The mean of these splines was taken across all subjects (since no significant MM-age relationships were observed before (Hui et al., 2022b) to generate the cohort-mean MM basis function. Water reference spectra were modeled with a simulated water basis function in the frequency domain with a 6-parameter model (amplitude, zero- and first-order phase, Gaussian and Lorentzian line-broadening, and frequency shift). Water-referenced metabolite concentrations were calculated according to (Gasparovic et al., 2006), adjusted for tissue-specific water visibility and relaxation times based on literature values (Wansapura et al., 1999) for each segmented tissue fraction of the voxel. Signal-to-noise ratio (SNR) was determined as the ratio of the maximum amplitude of the tNAA signal divided by the standard deviation of the noise, estimated from a de-trended signal-free area of the spectrum. The full-width at half-maximum (FWHM) linewidth of the tNAA signal was also determined.

### Statistical Analysis

All statistical analyses were performed using R (Version 4.1.1) in RStudio (Version 1.2.5019, Integrated Development for R. RStudio, PBC, Boston, MA). Data were analyzed for total NAA (tNAA = NAA+NAAG); total choline (tCho = GPC + PCh); total creatine (tCr = Cr + PCr); Glx = Glu + Gln; and individual contributions from GABA; Gln; Glu; GSH; mI; Lac; NAA; NAAG; PE; sI. Concentrations equal to 0 were interpreted as evidence of failure to fit and those datapoints were excluded from further analysis. Separate Pearson correlations were run assessing relationship between age and metabolite concentration in each voxel. Inter-metabolite Pearson correlations were also generated. Paired t-tests were used to investigate regional within-subject concentration differences between the PCC and CSO regions.

## Results

Data were excluded for one subject due to the detection of ethanol signals in the spectra. Data from one further subject were excluded in each region due to the appearance of unacceptably large lipid signals, presumably due to subject motion. The average NAA SNR was 160 in CSO and 150 in PCC, and the average NAA linewidth was 6.8 Hz in CSO and 7.0 Hz in PCC, as shown in Figure 2, indicating high data quality consistent with the acquisition parameters and the relatively favorable voxel locations.

**Figure 2.**
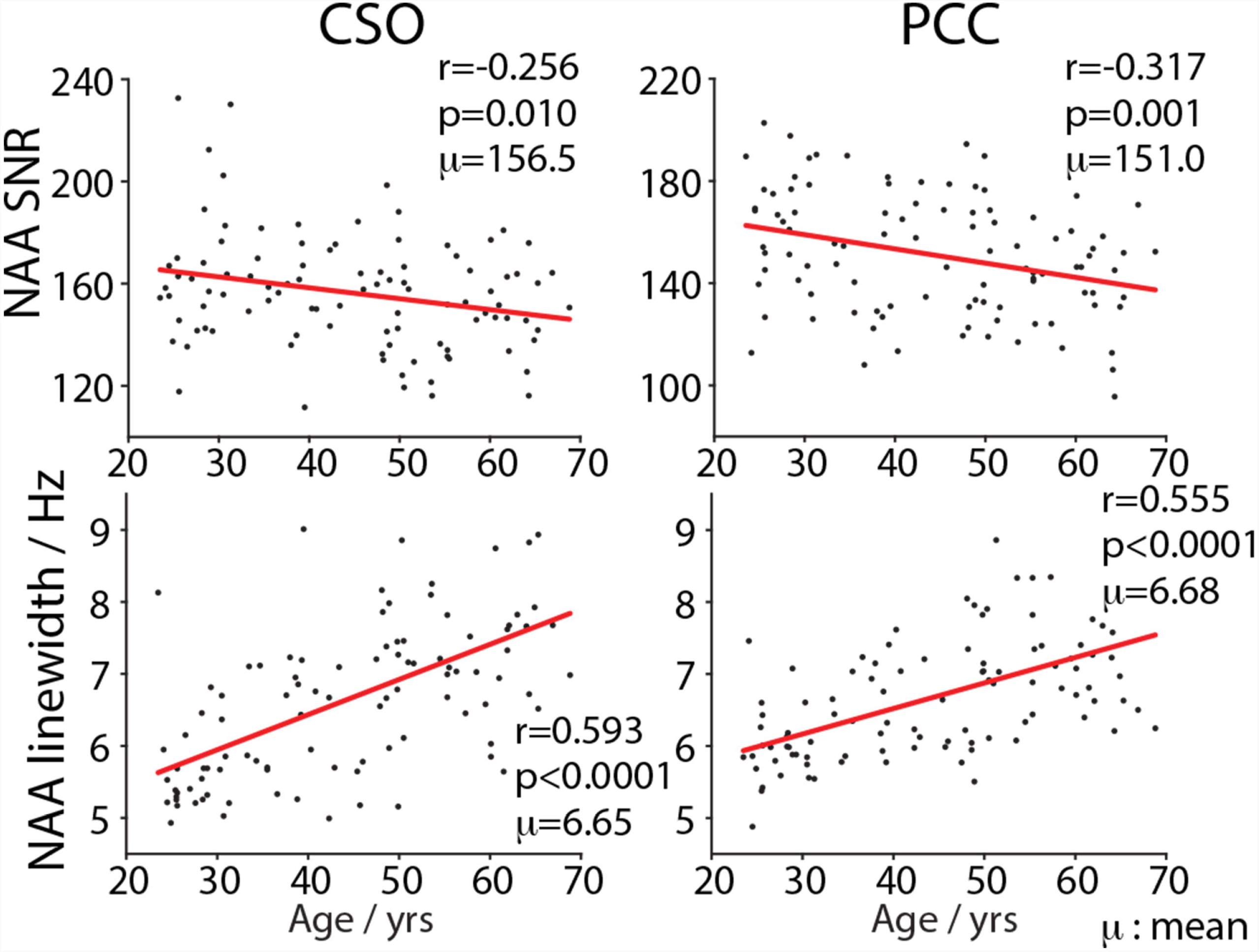
Correlations of age with the SNR and linewidth metrics of NAA. μ= mean.

Levels of Asp (t=-18, p<0.0001), GABA (t=-10, p<0.0001), Glx (t=-30, p<0.0001), GSH (t=-16, p<0.0001), Lac (t=-3.8, p<0.001), mI (t=-38, p<0.0001), sI (t=-14, p<0.0001), and tCr (t=-44, p<0.0001) were significantly higher in PCC, while tCho (t=27, p<0.0001) and tNAA (t=6.4, p<0.0001) were significantly higher in CSO.

Average spectra per decade from each region are shown in Figure 3. Age-by-metabolite correlation plots for the major metabolites are seen in Figure 4. Statistically significant age-by-metabolite correlations were seen for tCr (R^2^=0.36; p=0.0003), tCho (R^2^=0.11; p<0.001), sI (R^2^=0.11; p=0.004), and mI (R^2^=0.10; p<0.0003) in CSO, and tCr (R^2^=0.15; p<0.0001), tCho (R^2^=0.11; p<0.001), and GABA (R^2^=0.11; p=0.003) in PCC. No significant correlations were seen between tNAA, NAA, Glx or Glu and age in either region (all p>0.25). There is also significant positive correlation between linewidth and age, as seen in Figure 2, with older subjects tending to have broader signals. This is also reflected in SNR (peak height being inversely related to linewidth for a given area), as expected. Age correlations and inter-metabolite correlations are illustrated in Figure 5.

**Figure 3.**
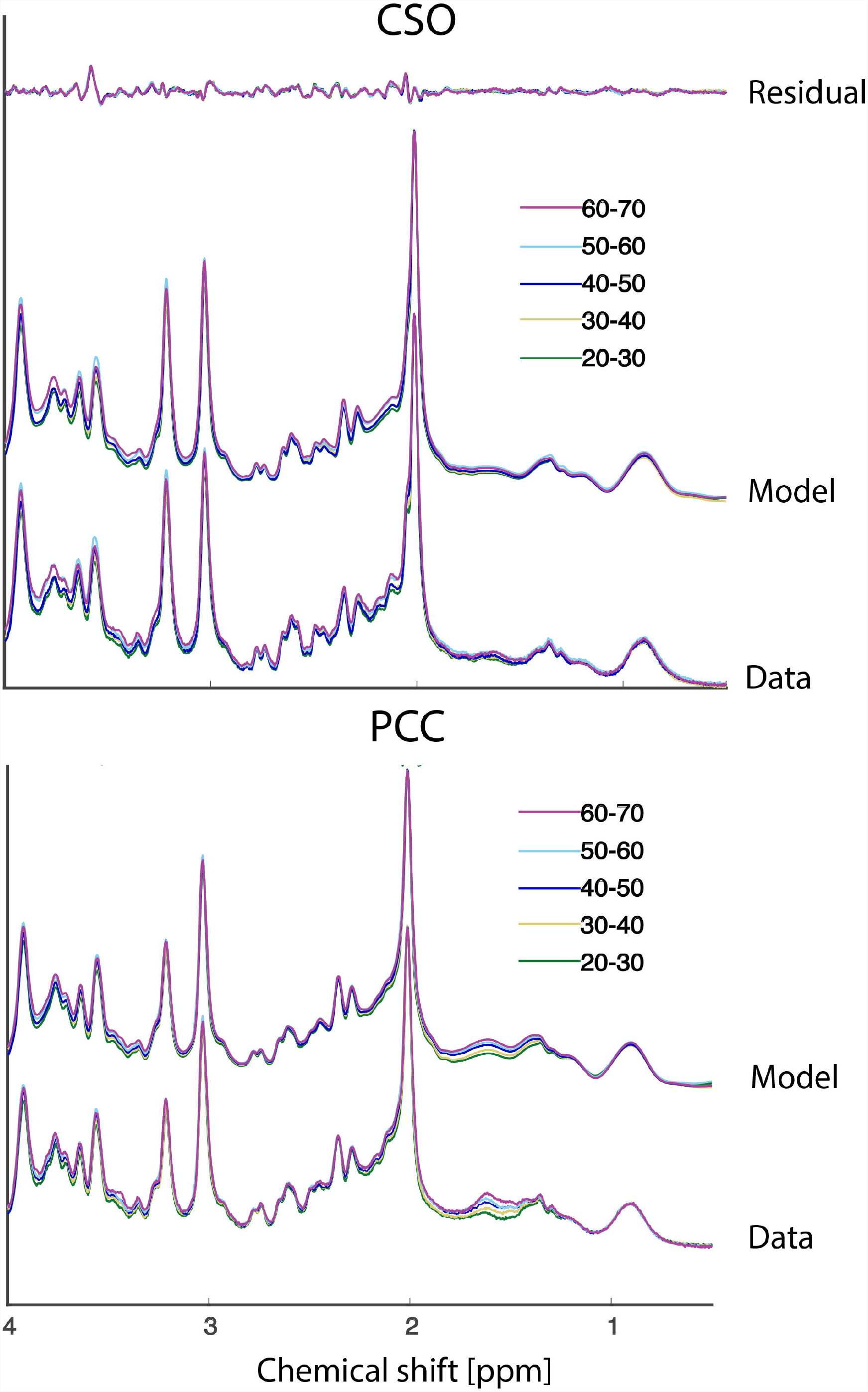
Average spectra per decade from CSO (upper panel) and PCC (lower panel).

**Figure 4.**
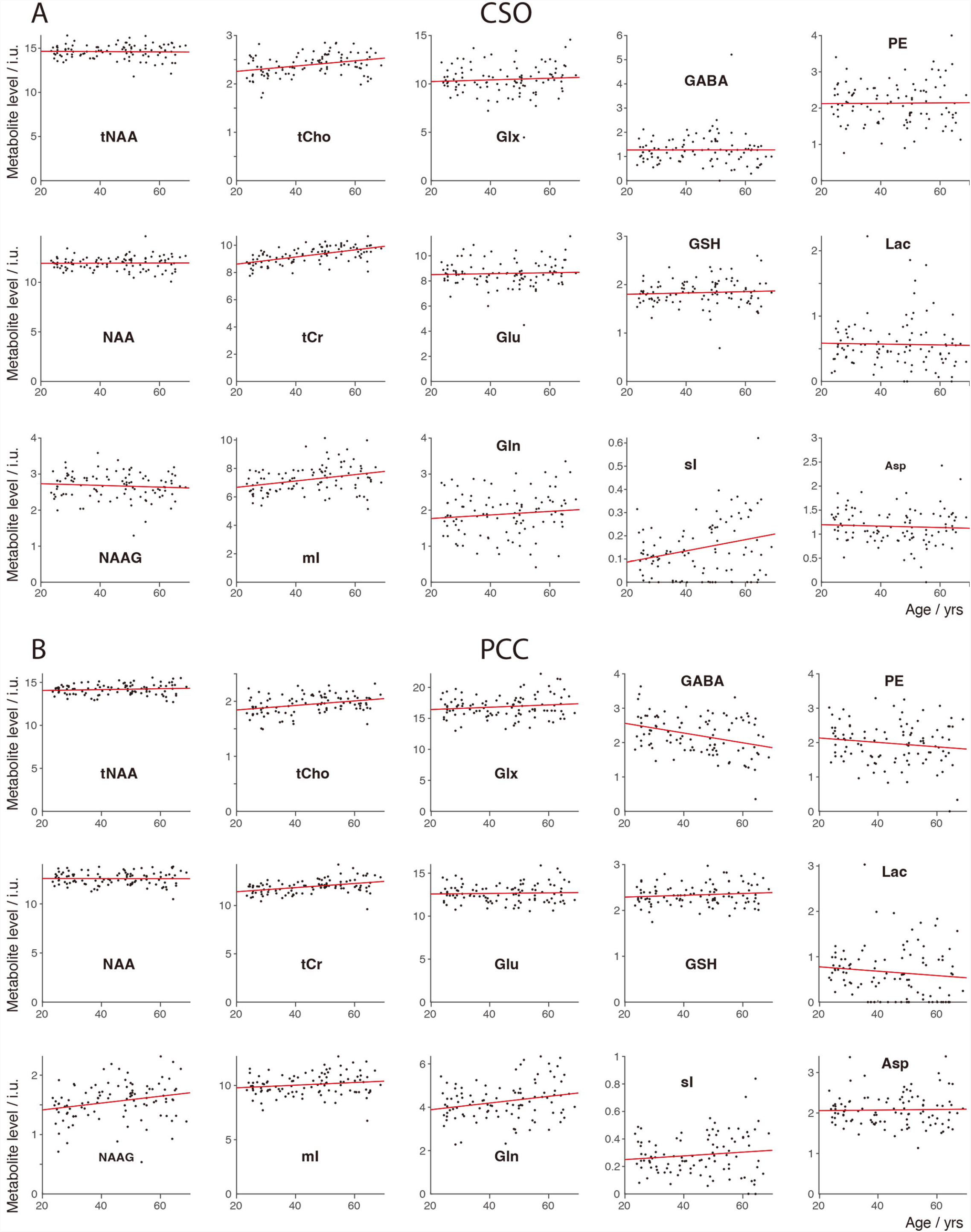
Metabolite-age correlation plots from (a) CSO and (b) PCC.

**Figure 5.**
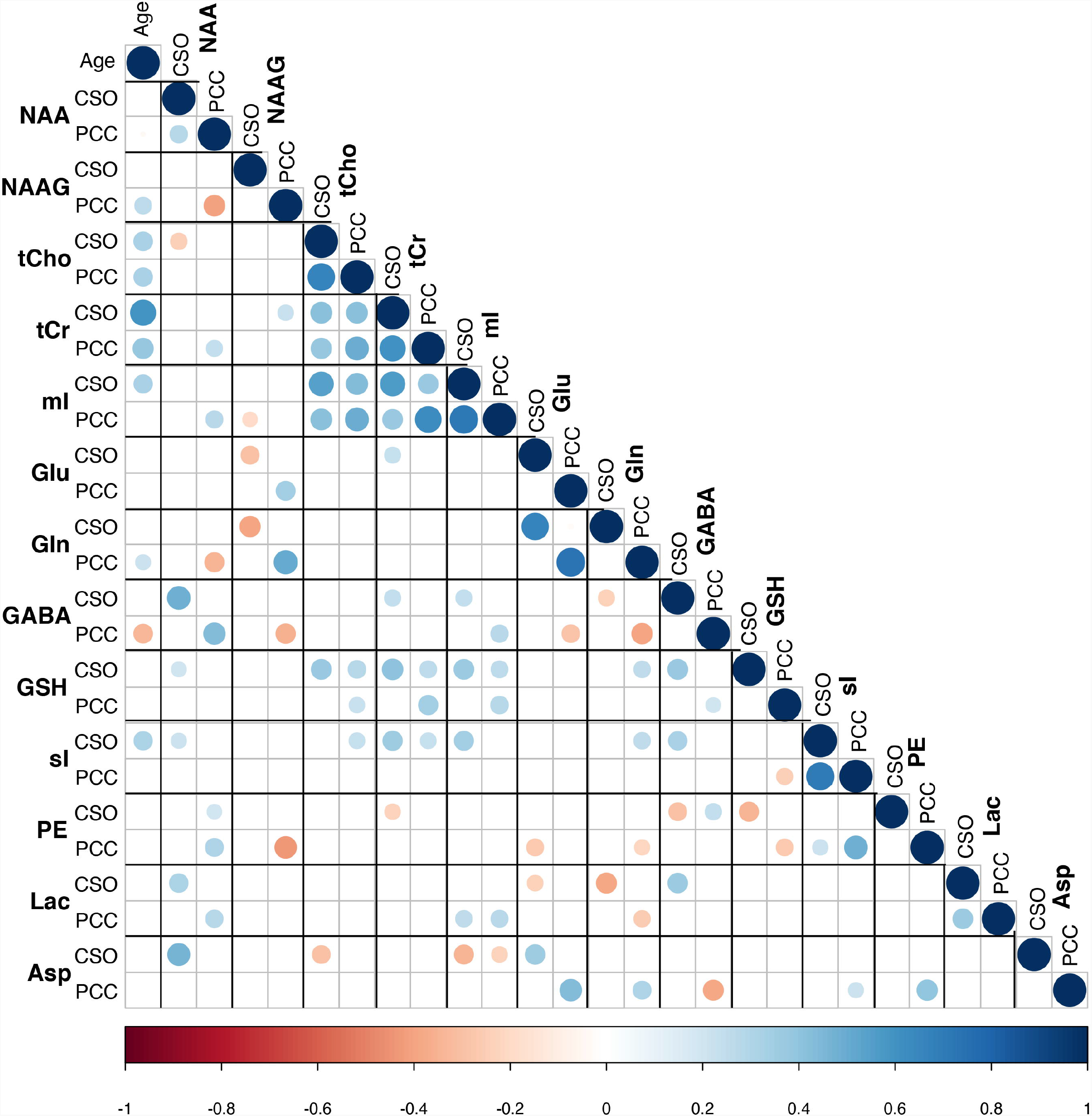
Age correlations and inter-metabolite correlations. Dot radius and color indicate correlation strength, while color indicates correlation directionality.

## Discussion

In this study, MRS data from a large structured cross-sectional cohort of male and female subjects throughout adulthood were investigated for neurometabolic changes as a function of age, using MRS consensus-recommended quantification methods (Near et al., 2021; Wilson et al., 2019). Positive age correlations in tCho, and tCr were observed for CSO and PCC, while none were found for NAA, tNAA, Glu or Glx in either region. The study further found GABA level decreased with age in PCC, increased sI levels in CSO, and significantly higher sI in the PCC for female subjects. Our results provide a normative assessment of the trajectories of MRS-measured metabolite levels in CSO and PCC across the healthy adult lifespan.

This study indicated that tCho and tCr increased with age in both CSO and PCC, in line with most previous MRS studies (Cleeland et al., 2019), perhaps driven by glial proliferation, as higher Cho and Cr levels are found in glial cells (Brand et al., 1993). The glial metabolite mI demonstrated a positive correlation with age in CSO, and a trend toward increase with age in PCC, that did not reach statistical significance. The metabolically linked sI showed a similar pattern (increase in CSO, no effect in PCC), in line with prior work (Kaiser et al., 2005). sI was also significantly higher in the PCC in female subjects, suggesting sex should be considered when investigating sI alterations.

The relatively strong positive correlations between tCr levels and age suggest that more age-related declines in metabolite levels would be reported if Cr-referenced metabolite ratios were used for quantification. While the quantification approach used here (relaxation and tissue correction based on literature reference values) complies with community consensus (Near et al., 2021; Wilson et al., 2019), it is still not free from potential confounds. There is strong literature evidence of age-related changes in the relaxation rates of water signals (Knight et al., 2016; Söderberg et al., 1990) and metabolite signals (Deelchand et al., 2020; Kirov et al., 2008; McIntyre et al., 2007; Schenker et al., 1993), that are not considered by the quantification approach used here. There is a strong need in the community for age-normed reference values to address this deficiency.

It is notable that age correlations were not observed for NAA, tNAA, Glu or Glx in either region, in spite of the fact that this is one of the commonest findings in the literature (Cleeland et al., 2019). One potential explanation for this is the age-range of our cohort (20-69 years), which would not be sensitive to changes later in life. NAA and Glu declines often seen in metabolite ratios – most often ratios to Cr – and so might be driven by the reference denominator as much as changes in the numerator. It may also be the case that non-linear metabolite by age relationships will be revealed in studies of wider age ranges, as we recently reported in a meta-analysis of edited GABA MRS across the lifespan (Ec et al., 2021). The majority of studies applying relaxation and tissue correction do not show age-related NAA changes (Wu et al., 2012).

However, a recent study applying absolute quantification by phantom (Kirov et al., 2021) did report decreases in whole-brain NAA concentration with age. This cohort included a large number of subjects above the age range of our cohort, so the difference in findings could be driven by either cohort, regional variation or methodology.

It is interesting to note that, in spite of short-TE PRESS being a somewhat controversial approach to measuring GABA levels, these data demonstrated significantly higher levels of GABA in GM than WM, and a reduction in GABA level with age. Both of these results are commonly seen in the edited GABA literature (Porges et al., 2017). The potential linewidth confound is also noteworthy – with increasing linewidth, modeling will find it increasingly difficult to resolve GABA signals from those of Glu, Gln and macromolecules. Indeed, estimates of GABA from linear-combination modeling do depend on SNR and increased linewidth (Near et al., 2013). Contrastingly, no age-related changes in GSH were observed, in contrast to the edited MRS literature (Hupfeld et al., 2021).

The inter-metabolite correlations reveal some interesting relationships. Overall, there are more positive than negative correlations. While there is a concern that common variance in the reference signal drive such correlations, they do not appear to be more prevalent within-region (common reference) compared to between region (independent reference). We therefore interpret this as reflecting genuine biological covariance. Several metabolites showed substantial positive correlations between PCC and CSO - tNAA, tCho, tCr, mI, and sI. Negative correlations between metabolites with overlapping basis spectra are potential evidence of the limitation of linear-combination modeling at 3T. These are seen between GABA and Glx, and for GSH and PE, both exclusively within-region.

We used the publicly available Osprey algorithm for perform linear-combination modeling of our data. While similar in concept, implementation and performance to the de-facto gold standard LCModel and other widely used methods like Tarquin (Wilson et al., 2011), we have recently demonstrated that results obtained with different modeling algorithms might differ in systematic fashion (Zöllner et al., 2021), a phenomenon commonly encountered in many neuroimaging disciplines. One key algorithmic difference is that the Osprey algorithm does not apply soft constraints to regularize the contributions from typically low-concentration metabolites like GABA and GSH, as is done by the LCModel, which may decrease systematic biases.

There are some limitations in this study, first, even though an adequate sample size was achieved, the age span was relatively narrow. In particular, we did not enroll participants above 70 years of age, i.e. when effects of aging drastically accelerate. Future MRS-aging studies should increase the age range to establish normative age trajectories during this important late-life stage. Second, only two selected ROIs (PCC and CSO) were analyzed in this study, one is gray-matter predominant region, another white-matter predominant. Neurochemical changes during aging are highly region-dependent (Eylers et al., 2016), and data from more regions or even whole brain data will be needed to improve our understanding of age-related changes.

## Conclusion

The results indicated positive age correlations in tCho, tCr, and mI for CSO and in tCho, tCr, and GABA for PCC, while no age-related changes for NAA, tNAA, Glu or Glx. Our results provide further evidence of neurometabolic time course of healthy aging, suggesting that age matching is essential for comparative studies of neuro-degenerative disease using MRS.

## Acknowledgements

This work was supported by Natural Science Foundation of Shandong (grant number: ZR2020QH267); Major Research Project of Shandong Province (grant number: 2016ZDJS07A16); and National Institutes of Health (grant number: R01 EB016089; P41 EB031771; R00 AG062230; K00 AG068440).

## Disclosures of Conflicts of Interest

All authors declare no conflicts of interest.

## Notes

### Competing Interest Statement

The authors have declared no competing interest.

## References

Brand, A., Richter-Landsberg, C., Leibfritz, D., 1993. Multinuclear NMR Studies on the Energy Metabolism of Glial and Neuronal Cells. DNE 15, 289–298. https://doi.org/10.1159/000111347

Cheng, H., Wang, A., Newman, S., Dydak, U., 2021. An investigation of glutamate quantification with PRESS and MEGA-PRESS. NMR in Biomedicine 34, e4453. https://doi.org/10.1002/nbm.4453

Cleeland, C., Pipingas, A., Scholey, A., White, D., 2019. Neurochemical changes in the aging brain: A systematic review. Neurosci Biobehav Rev 98, 306–319. https://doi.org/10.1016/j.neubiorev.2019.01.003

Dai, G., Yu, H., Kruse, M., Traynor-Kaplan, A., Hille, B., 2016. Osmoregulatory inositol transporter SMIT1 modulates electrical activity by adjusting PI(4,5)P2 levels. Proc Natl Acad Sci U S A 113, E3290–3299. https://doi.org/10.1073/pnas.1606348113

Deelchand, D.K., McCarten, J.R., Hemmy, L.S., Auerbach, E.J., Eberly, L.E., Marjańska, M., 2020. Changes in the intracellular microenvironment in the aging human brain. Neurobiology of Aging 95, 168–175. https://doi.org/10.1016/j.neurobiolaging.2020.07.017

Diedenhofen, B., Musch, J., 2015. cocor: A Comprehensive Solution for the Statistical Comparison of Correlations. PLOS ONE 10, e0121945. https://doi.org/10.1371/journal.pone.0121945

Dwivedi, D., Megha, K., Mishra, R., Mandal, P.K., 2020. Glutathione in Brain: Overview of Its Conformations, Functions, Biochemical Characteristics, Quantitation and Potential Therapeutic Role in Brain Disorders. Neurochem Res 45, 1461–1480. https://doi.org/10.1007/s11064-020-03030-1

Ec, P., g, J., b, F., Ra, E., Na, P., 2021. The trajectory of cortical GABA across the lifespan, an individual participant data meta-analysis of edited MRS studies. eLife 10. https://doi.org/10.7554/eLife.62575

Eylers, V.V., Maudsley, A.A., Bronzlik, P., Dellani, P.R., Lanfermann, H., Ding, X.-Q., 2016. Detection of Normal Aging Effects on Human Brain Metabolite Concentrations and Microstructure with Whole-Brain MR Spectroscopic Imaging and Quantitative MR Imaging. American Journal of Neuroradiology 37, 447–454. https://doi.org/10.3174/ajnr.A4557

Friston, K.J., Holmes, A.P., Worsley, K.J., Poline, J.-P., Frith, C.D., Frackowiak, R.S.J., 1994. Statistical parametric maps in functional imaging: A general linear approach. Human Brain Mapping 2, 189–210. https://doi.org/10.1002/hbm.460020402

Gao, F., Edden, R.A.E., Li, M., Puts, N.A.J., Wang, G., Liu, C., Zhao, B., Wang, H., Bai, X., Zhao, C., Wang, X., Barker, P.B., 2013. Edited magnetic resonance spectroscopy detects an age-related decline in brain GABA levels. Neuroimage 78, 75–82. https://doi.org/10.1016/j.neuroimage.2013.04.012

Gasparovic, C., Song, T., Devier, D., Bockholt, H.J., Caprihan, A., Mullins, P.G., Posse, S., Jung, R.E., Morrison, L.A., 2006. Use of tissue water as a concentration reference for proton spectroscopic imaging. Magn Reson Med 55, 1219–1226. https://doi.org/10.1002/mrm.20901

Haga, K.K., Khor, Y.P., Farrall, A., Wardlaw, J.M., 2009. A systematic review of brain metabolite changes, measured with 1H magnetic resonance spectroscopy, in healthy aging. Neurobiol Aging 30, 353–363. https://doi.org/10.1016/j.neurobiolaging.2007.07.005

Harris, A.D., Saleh, M.G., Edden, R.A.E., 2017. Edited 1H magnetic resonance spectroscopy in vivo: Methods and metabolites. Magnetic Resonance in Medicine 77, 1377–1389. https://doi.org/10.1002/mrm.26619

Howe, F.A., Barton, S.J., Cudlip, S.A., Stubbs, M., Saunders, D.E., Murphy, M., Wilkins, P., Opstad, K.S., Doyle, V.L., McLean, M.A., Bell, B.A., Griffiths, J.R., 2003. Metabolic profiles of human brain tumors using quantitative in vivo1H magnetic resonance spectroscopy. Magn. Reson. Med. 49, 223–232. https://doi.org/10.1002/mrm.10367

Hoyer, C., Gass, N., Weber-Fahr, W., Sartorius, A., 2014. Advantages and Challenges of Small Animal Magnetic Resonance Imaging as a Translational Tool. NPS 69, 187–201. https://doi.org/10.1159/000360859

Hui, S.C.N., Gong, T., Zöllner, H.J., Song, Y., Murali-Manohar, S., Oeltzschner, G., Mikkelsen, M., Tapper, S., Chen, Y., Saleh, M.G., Porges, E.C., Chen, W., Wang, G., Edden, R.A.E., 2022a. The macromolecular MR spectrum does not change with healthy aging. Magn Reson Med 87, 1711–1719. https://doi.org/10.1002/mrm.29093

Hui, S.C.N., Gong, T., Zöllner, H.J., Song, Y., Murali-Manohar, S., Oeltzschner, G., Mikkelsen, M., Tapper, S., Chen, Y., Saleh, M.G., Porges, E.C., Chen, W., Wang, G., Edden, R.A.E., 2022b. The macromolecular MR spectrum does not change with healthy aging. Magnetic Resonance in Medicine 87, 1711–1719. https://doi.org/10.1002/mrm.29093

Hupfeld, K.E., Hyatt, H.W., Alvarez Jerez, P., Mikkelsen, M., Hass, C.J., Edden, R.A.E., Seidler, R.D., Porges, E.C., 2021. In Vivo Brain Glutathione is Higher in Older Age and Correlates with Mobility. Cerebral Cortex 31, 4576–4594. https://doi.org/10.1093/cercor/bhab107

Kaiser, L.G., Schuff, N., Cashdollar, N., Weiner, M.W., 2005. Scyllo-inositol in normal aging human brain: 1H magnetic resonance spectroscopy study at 4 Tesla. NMR Biomed 18, 51–55. https://doi.org/10.1002/nbm.927

Kirov, I.I., Fleysher, L., Fleysher, R., Patil, V., Liu, S., Gonen, O., 2008. The Age Dependence of Regional Proton Metabolites T2 Relaxation Times in the Human Brain at 3 Tesla. Magn Reson Med 60, 790–795. https://doi.org/10.1002/mrm.21715

Kirov, I.I., Sollberger, M., Davitz, M.S., Glodzik, L., Soher, B.J., Babb, J.S., Monsch, A.U., Gass, A., Gonen, O., 2021. Global brain volume and N-acetyl-aspartate decline over seven decades of normal aging. Neurobiol Aging 98, 42–51. https://doi.org/10.1016/j.neurobiolaging.2020.10.024

Knight, M.J., McCann, B., Tsivos, D., Couthard, E., Kauppinen, R.A., 2016. Quantitative T1 and T2 MRI signal characteristics in the human brain: different patterns of MR contrasts in normal ageing. MAGMA 29, 833–842. https://doi.org/10.1007/s10334-016-0573-0

Landim, R.C.G., Edden, R.A.E., Foerster, B., Li, L.M., Covolan, R.J.M., Castellano, G., 2016. Investigation of NAA and NAAG dynamics underlying visual stimulation using MEGA-PRESS in a functional MRS experiment. Magn Reson Imaging 34, 239–245. https://doi.org/10.1016/j.mri.2015.10.038

Langa, K.M., Larson, E.B., Wallace, R.B., Fendrick, A.M., Foster, N.L., Kabeto, M.U., Weir, D.R., Willis, R.J., Herzog, A.R., 2004. Out-of-pocket health care expenditures among older Americans with dementia. Alzheimer Disease and Associated Disorders 18, 90–98. https://doi.org/10.1097/01.wad.0000126620.73791.3e

Maes, C., Hermans, L., Pauwels, L., Chalavi, S., Leunissen, I., Levin, O., Cuypers, K., Peeters, R., Sunaert, S., Mantini, D., Puts, N.A.J., Edden, R.A.E., Swinnen, S.P., 2018. Age-related differences in GABA levels are driven by bulk tissue changes. Human Brain Mapping 39, 3652–3662. https://doi.org/10.1002/hbm.24201

McIntyre, D.J.O., Charlton, R.A., Markus, H.S., Howe, F.A., 2007. Long and short echo time proton magnetic resonance spectroscopic imaging of the healthy aging brain. Journal of Magnetic Resonance Imaging 26, 1596–1606. https://doi.org/10.1002/jmri.21198

McLaurin, J., Golomb, R., Jurewicz, A., Antel, J.P., Fraser, P.E., 2000. Inositol Stereoisomers Stabilize an Oligomeric Aggregate of Alzheimer Amyloid β Peptide and Inhibit Aβ-induced Toxicity *. Journal of Biological Chemistry 275, 18495–18502. https://doi.org/10.1074/jbc.M906994199

Menshchikov, P.E., Akhadov, T.A., Semenova, N.A., 2017. Quantification of cerebral aspartate concentration in vivo using proton magnetic resonance spectroscopy. Bull. Lebedev Phys. Inst. 44, 56–60. https://doi.org/10.3103/S1068335617030022

Morana, G., Piccardo, A., Puntoni, M., Nozza, P., Cama, A., Raso, A., Mascelli, S., Massollo, M., Milanaccio, C., Garrè, M.L., Rossi, A., 2015. Diagnostic and prognostic value of 18F-DOPA PET and 1H-MR spectroscopy in pediatric supratentorial infiltrative gliomas: a comparative study. Neuro-Oncology 17, 1637–1647. https://doi.org/10.1093/neuonc/nov099

Mullins, P.G., McGonigle, D.J., O’Gorman, R.L., Puts, N.A.J., Vidyasagar, R., Evans, C.J., Cardiff Symposium on MRS of GABA, Edden, R.A.E., 2014. Current practice in the use of MEGA-PRESS spectroscopy for the detection of GABA. Neuroimage 86, 43–52. https://doi.org/10.1016/j.neuroimage.2012.12.004

Near, J., Andersson, J., Maron, E., Mekle, R., Gruetter, R., Cowen, P., Jezzard, P., 2013. Unedited in vivo detection and quantification of γ-aminobutyric acid in the occipital cortex using short-TE MRS at 3 T. NMR in Biomedicine 26, 1353–1362. https://doi.org/10.1002/nbm.2960

Near, J., Harris, A.D., Juchem, C., Kreis, R., Marjańska, M., Öz, G., Slotboom, J., Wilson, M., Gasparovic, C., 2021. Preprocessing, analysis and quantification in single-voxel magnetic resonance spectroscopy: experts’ consensus recommendations. NMR Biomed 34, e4257. https://doi.org/10.1002/nbm.4257

Oeltzschner, G., Zöllner, H.J., Hui, S.C.N., Mikkelsen, M., Saleh, M.G., Tapper, S., Edden, R.A.E., 2020. Osprey: Open-source processing, reconstruction & estimation of magnetic resonance spectroscopy data. Journal of Neuroscience Methods 343, 108827. https://doi.org/10.1016/j.jneumeth.2020.108827

Porges, E.C., Jensen, G., Foster, B., Edden, R.A., Puts, N.A., 2021. The trajectory of cortical GABA across the lifespan, an individual participant data meta-analysis of edited MRS studies. eLife 10, e62575. https://doi.org/10.7554/eLife.62575

Porges, E.C., Woods, A.J., Lamb, D.G., Williamson, J.B., Cohen, R.A., Edden, R.A.E., Harris, A.D., 2017. Impact of tissue correction strategy on GABA-edited MRS findings. NeuroImage 162, 249–256. https://doi.org/10.1016/j.neuroimage.2017.08.073

Ross, A.J., Sachdev, P.S., 2004. Magnetic resonance spectroscopy in cognitive research. Brain Research Reviews 44, 83–102. https://doi.org/10.1016/j.brainresrev.2003.11.001

Schenker, C., Meier, D., Wichmann, W., Boesiger, P., Valavanis, A., 1993. Age distribution and iron dependency of the T2 relaxation time in the globus pallidus and putamen. Neuroradiology 35, 119–124. https://doi.org/10.1007/BF00593967

Simpson, R., Devenyi, G.A., Jezzard, P., Hennessy, T.J., Near, J., 2017. Advanced processing and simulation of MRS data using the FID appliance (FID-A)-An open source, MATLAB-based toolkit. Magn Reson Med 77, 23–33. https://doi.org/10.1002/mrm.26091

Söderberg, M., Edlund, C., Kristensson, K., Dallner, G., 1990. Lipid Compositions of Different Regions of the Human Brain During Aging. Journal of Neurochemistry 54, 415–423. https://doi.org/10.1111/j.1471-4159.1990.tb01889.x

Tkác, I., Starcuk, Z., Choi, I.Y., Gruetter, R., 1999. In vivo 1H NMR spectroscopy of rat brain at 1 ms echo time. Magn Reson Med 41, 649–656. https://doi.org/10.1002/(sici)1522-2594(199904)41:4<649::aid-mrm2>3.0.co;2-g

Wansapura, J.P., Holland, S.K., Dunn, R.S., Ball Jr., W.S., 1999. NMR relaxation times in the human brain at 3.0 tesla. Journal of Magnetic Resonance Imaging 9, 531–538. https://doi.org/10.1002/(SICI)1522-2586(199904)9:4<531::AID-JMRI4>3.0.CO;2-L

Wilson, M., Andronesi, O., Barker, P.B., Bartha, R., Bizzi, A., Bolan, P.J., Brindle, K.M., Choi, I.-Y., Cudalbu, C., Dydak, U., Emir, U.E., Gonzalez, R.G., Gruber, S., Gruetter, R., Gupta, R.K., Heerschap, A., Henning, A., Hetherington, H.P., Huppi, P.S., Hurd, R.E., Kantarci, K., Kauppinen, R.A., Klomp, D.W.J., Kreis, R., Kruiskamp, M.J., Leach, M.O., Lin, A.P., Luijten, P.R., Marjańska, M., Maudsley, A.A., Meyerhoff, D.J., Mountford, C.E., Mullins, P.G., Murdoch, J.B., Nelson, S.J., Noeske, R., Öz, G., Pan, J.W., Peet, A.C., Poptani, H., Posse, S., Ratai, E.-M., Salibi, N., Scheenen, T.W.J., Smith, I.C.P., Soher, B.J., Tkáč, I., Vigneron, D.B., Howe, F.A., 2019. Methodological consensus on clinical proton MRS of the brain: Review and recommendations. Magn Reson Med 82, 527–550. https://doi.org/10.1002/mrm.27742

Wilson, M., Reynolds, G., Kauppinen, R.A., Arvanitis, T.N., Peet, A.C., 2011. A constrained least-squares approach to the automated quantitation of in vivo <sup>1</sup>H magnetic resonance spectroscopy data. Magn Reson Med 65, 1–12. https://doi.org/10.1002/mrm.22579

Wu, W.E., Gass, A., Glodzik, L., Babb, J.S., Hirsch, J., Sollberger, M., Achtnichts, L., Amann, M., Monsch, A.U., Gonen, O., 2012. Whole brain N-acetylaspartate concentration is conserved throughout normal aging. Neurobiol Aging 33, 2440–2447. https://doi.org/10.1016/j.neurobiolaging.2011.12.008

Zöllner, H.J., Považan, M., Hui, S.C.N., Tapper, S., Edden, R.A.E., Oeltzschner, G., 2021. Comparison of different linear-combination modeling algorithms for short-TE proton spectra. NMR Biomed 34, e4482. https://doi.org/10.1002/nbm.4482

